# Heavy Metal-Resistant, Plastic-Degrading *Bacillus* sp. Isolated from Landfill Leachate: Identification and Characterization

**DOI:** 10.64898/2026.02.27.708447

**Authors:** Umme Samia Antu, Amily Sarker, Nabila Haque, Joyoshrie Karmakar, Abdul Khaleque, Md. Sabir Hossain, Md. Anowar Khasru Parvez

## Abstract

Landfill leachates in rapidly urbanizing regions like Dhaka present a complex ecological challenge owing to the concurrent buildup of heavy metals and plastic waste. Despite the severity of this pollution, the role of indigenous multi-functional bacteria in mitigating these mixed contaminants remains poorly understood. This research sought to isolate and characterize bacteria resistant to heavy metals and capable of degrading plastics from the Aminbazar and Matuail landfills and evaluate their bioremediation potential. Physicochemical analysis confirmed extreme contamination, with heavy metal levels (Pb, Cr, Cd, Cu) significantly exceeding WHO safety limits. Out of 81 isolates, nearly half exhibited multi-metal resistance and polyethylene (PE) degradation capacity. Statistical analysis showed a significant correlation between plastic degradation and multi-metal tolerance, suggesting a linked evolutionary adaptation. Enzymatic assays confirmed enzymes (e.g., urease, catalase, citrate and esterase) as drivers of both plastic degradation and heavy metal tolerance in leading isolates. Molecular screening identified the resistance genes *pbrA* and *alkB*, while the high prevalence of Class 1 integrons (80% in *pbrA*-positive isolates) points to a high potential for horizontal gene transfer in these environments. Furthermore, MALDI-TOF MS identified the functional isolates as *Bacillus* sp. with FTIR verifying the contribution of specific cell-surface functional groups to metal biosorption. These results underscore the promise of native *Bacillus* strains as promising agents for the development of sustainable, integrated biotechnologies for landfill restoration and complex waste management.

## Introduction

Rapid industrialization and urbanization have markedly elevated global municipal solid waste (MSW) generation, which amounted to 2,010 million tons in 2016 and is projected to escalate to 3,400 million tons by 2050 [1]. Beyond the sheer volume of waste, its chemical toxicity represents a massive environmental burden; it is estimated that the global anthropogenic discharge of heavy metals into the ecosystem driven largely by the disposal of electronic and industrial waste in landfills includes approximately 1.2 billion kg of lead (Pb) and 3 million kg of cadmium (Cd) annually [2]; UNEP] [3]. Parallel to this chemical threat is the persistence of plastic waste, with global production reaching an estimated 8.3 billion tons to date, resulting in 4.9 billion tons accumulating in landfills or natural ecosystems [2]. In Bangladesh, the challenge is particularly acute; Dhaka alone produces 15,000 tons of waste daily, with only 37% reaching the Aminbazar and Matuail landfills [4–7]. Because landfills are the primary disposal method, they become concentrated sources of toxic leachate a complex wastewater containing heavy metals (HMs) and microplastics (MPs) that pose severe risks to public health and ecological integrity [5–8].

The environmental consequences of leachate are driven by the persistence of its chemical constituents. In Bangladesh, landfill leachate has been found to exceed environmental safety standards for metals such as Fe, Cu, and Ni [4–7]. Heavy metals, including Cd, Cr, and Pb, exert cytotoxic effects by disrupting essential microbial processes, such as protein synthesis and enzymatic activity, and causing oxidative DNA damage [9] [10] [11] [12]. Microplastics, primarily polyethylene terephthalate (PET), polyethylene (PE), and polypropylene (PP), were found at 270 ± 13 particles/L [13,14]. These particles act as vectors by adsorbing heavy metals, thereby forming synergistic pollutant complexes that are difficult to treat using conventional methods.

In response to this chronic exposure, indigenous microbial communities have evolved sophisticated resistance mechanisms. Microorganisms continuous exposure to heavy metals develop resistance mechanisms, including HMRGs (Heavy Metal Resistance genes) like *czcA* and *pbrA* [15], with bacteria such as *Bacillus subtilis, Pseudomonas aeruginosa* showing resistance to Hg, Cd, and Pb [16–18]. Significantly, recent metagenomic studies of landfill sites, such as those in Gujarat, India, have revealed that these microbiomes also harbor plastic-degrading genes [2].

In such polluted environments, HMRGs often co-occur with plastic-degrading genes on mobile genetic elements (MGEs) like Class 1 integrons. This co-selection, facilitated by horizontal gene transfer (HGT), allows certain genera including *Bacillus sp.*, *Acinetobacter sp.*, and *Pseudomonas sp.* to possess a dual repertoire for resistance and degradative traits [16,19,20]

Bioremediation emerges as a sustainable, cost-effective alternative to chemical detoxification by utilizing these microbial metabolic pathways [7] **(Fig 1)**. Microbial processes such as bioaccumulation, enzymatic transformation, and microbially induced calcite precipitation (MICP) via urease activity can effectively immobilize metals [21] [22–24]. Simultaneously, enzymes like esterases and nitrate reductases enhance the bioavailability and breakdown of organic pollutants [25].

**Figure 1:**
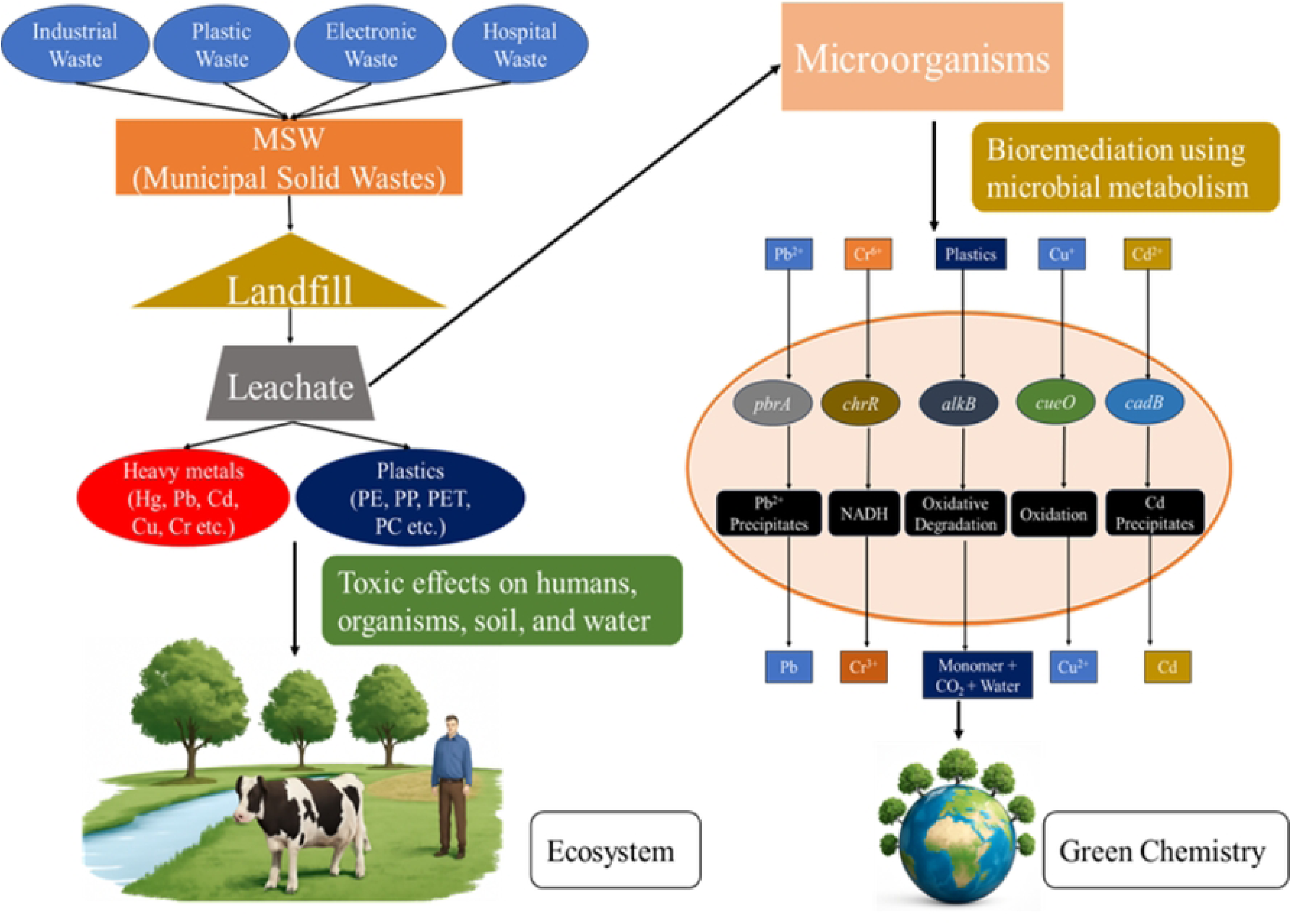
Bioremediation of Heavy Metals and Plastics in Landfill Leachate Using Microbial Metabolism.

Despite the promising potential of these indigenous microbes, a significant knowledge gap remains regarding the molecular characterization of multi-functional bacteria in Bangladesh. While the contamination levels of Dhaka’s landfills are documented, the specific bioremediation potential of local *Bacillus* strains concerning simultaneous heavy metal tolerance and plastic degradation has not been fully explored. Therefore, this research seeks to retrieve and characterize *Bacillus sp.* from landfill leachate and evaluate their capacity for metal removal and plastic degradation. This research represents an important step in addressing knowledge gaps in sustainable pollution management and provides a foundational report on the application of indigenous microbial isolates for environmental remediation in Bangladesh.

## Results

### Analysis of Physicochemical Parameters of Samples

A total 10 samples (8 Leachate and 2 Ground water sample) were collected to analyze physicochemical parameters. The temperature, electric conductivity (EC), total dissolved solids (TDS), salinity and dissolved oxygen (DO) profile of the leachate samples varied significantly from the ground water samples. In sample AB-1 and MT-1, TDS, EC, salinity are markedly high and DO which is markedly low **(Table 1)**. Based on these results, the raw leachate samples (AB-1 and MT-1) were prioritized for further investigation [5].

**Table 1.**
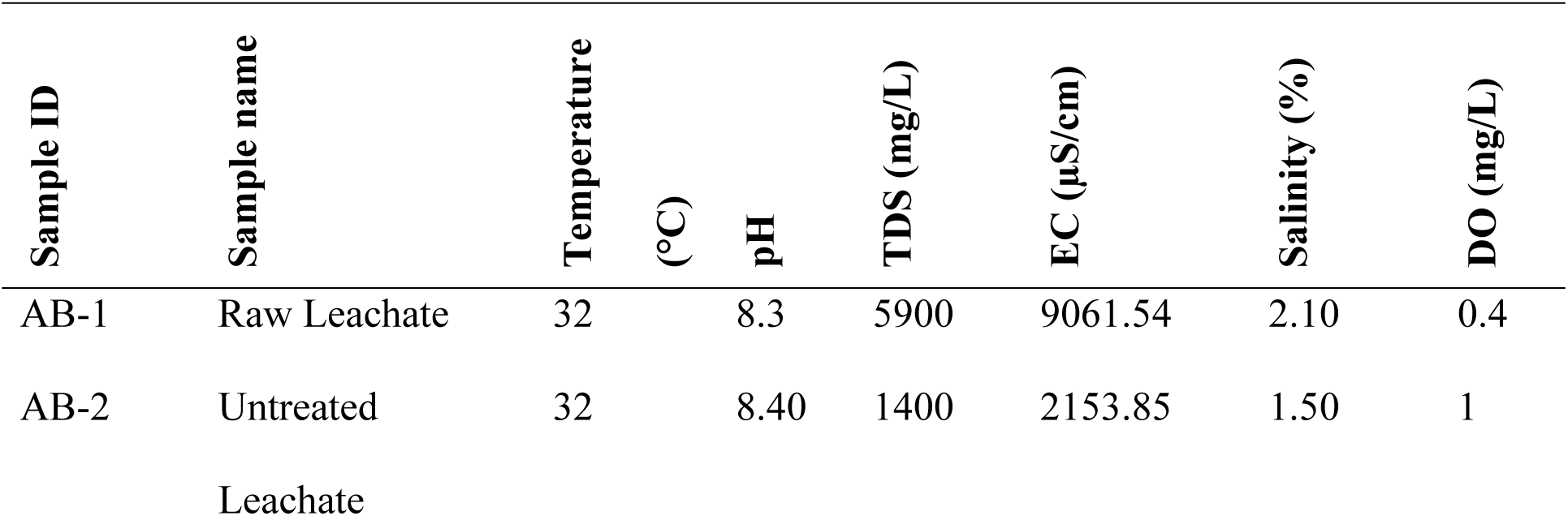
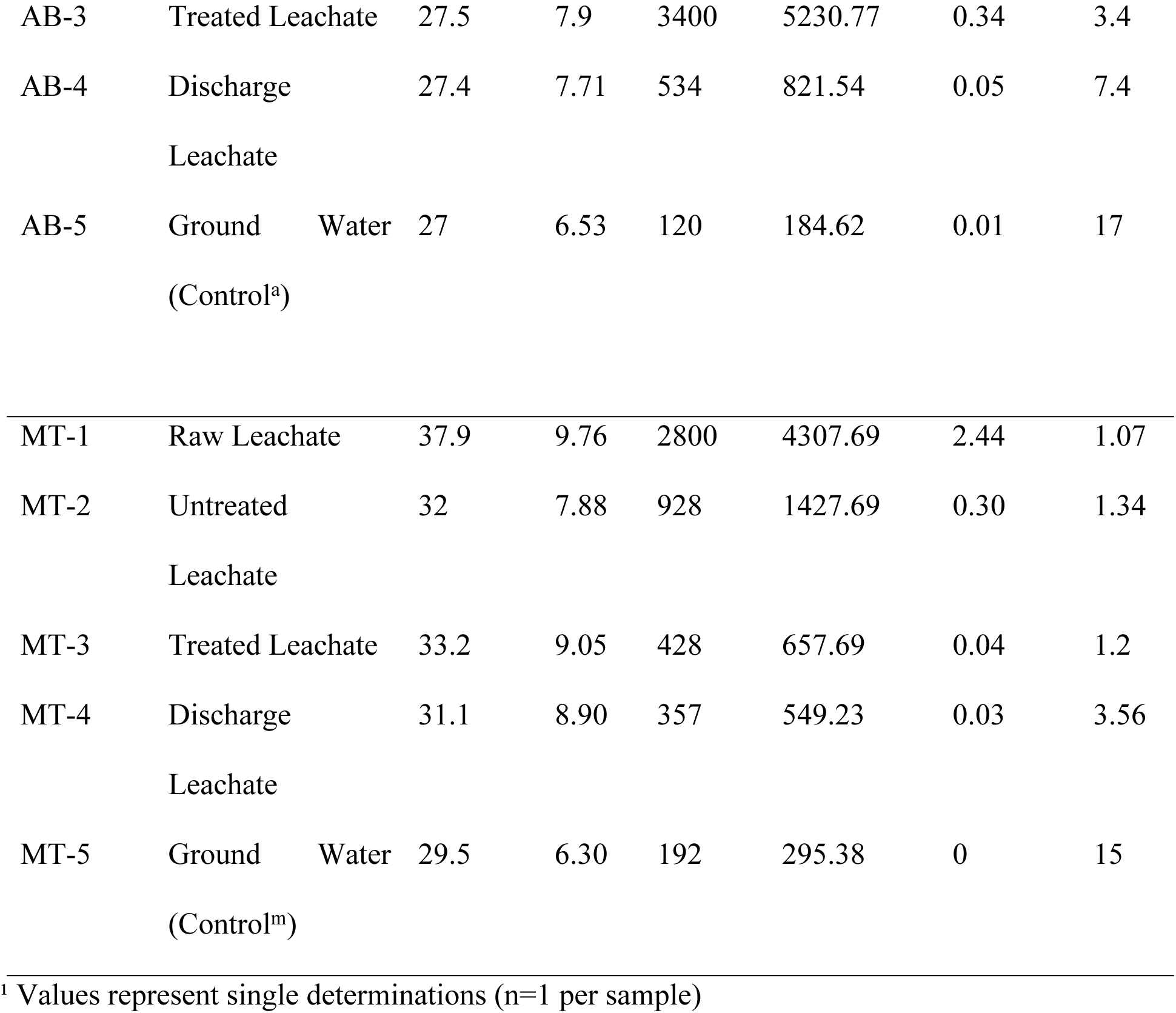
Physicochemical Parameters of the Landfill Leachate Samples ¹.

### Determination of Heavy-metal Concentration in Raw Leachate Samples

Heavy-metal concentration for Pb, Cr, Cd and Cu were examined from AB-1 and MT-1 by Atomic Absorption Spectroscopy (AAS). Which showed concentration variations of Pb (16.6 to 30.32 mg/l), Cr (14.06 to 24.58 mg/l), Cd (0.69 to 0.88 mg/l) along with Cu (5.47 to 21.05 mg/l) **(Fig 2 and S1 Table),** revealed that, all these heavy metal (Pb, Cr, Cd and Cu) concentrations exceeding the WHO permissible limits [5]. Additionally, the raw leachate of the Matuail landfill exhibits greater heavy metals concentration than that from the Aminbazar landfill [4].

**Fig 2:**
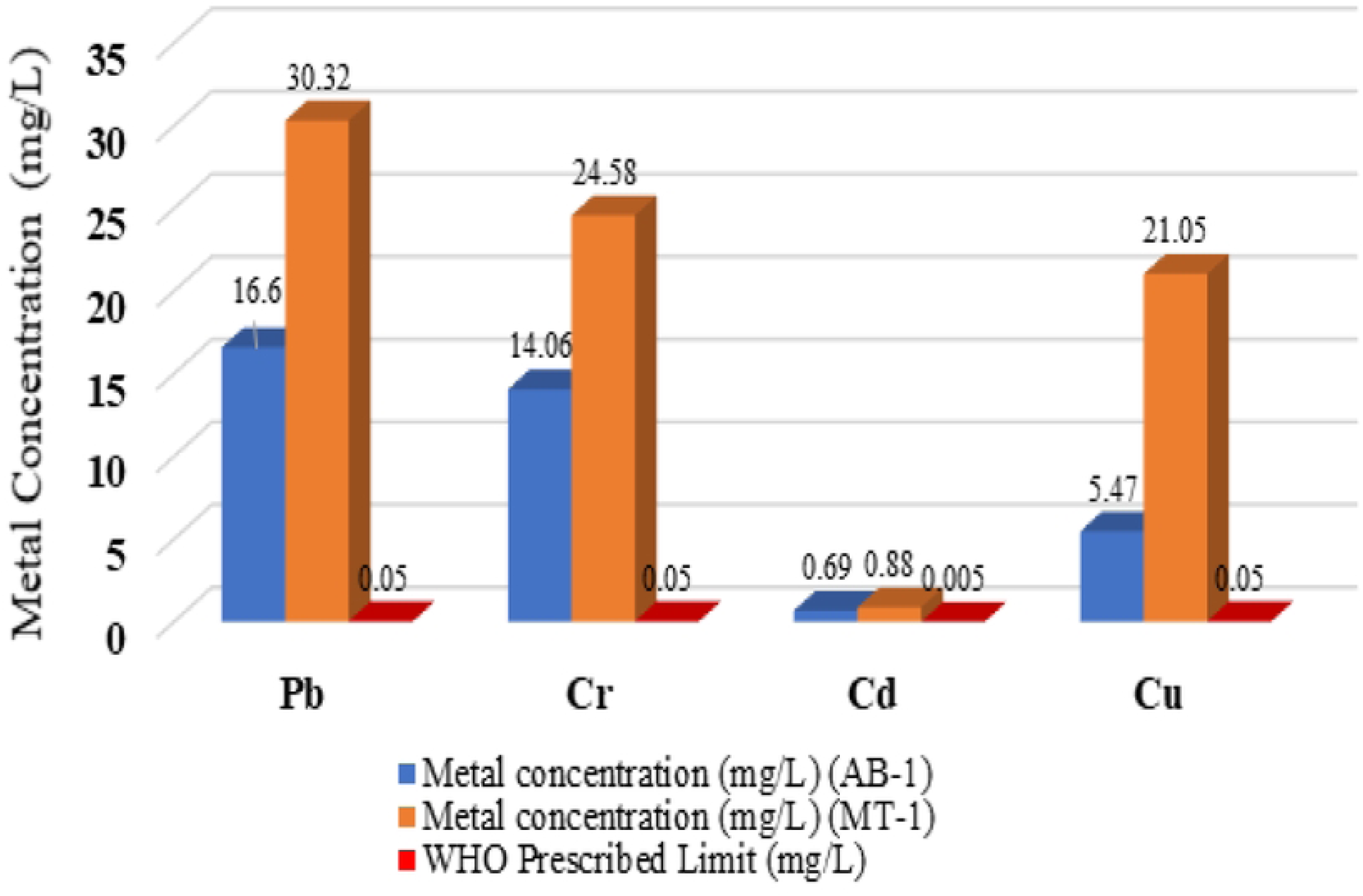
Heavy Metal Concentrations (mg/L) in Samples (AB-1 & MT-1) Compared to WHO Prescribed Limits.

### Enumeration of Total Heavy Metal-Resistant Bacteria and Isolation of Pure Colonies

Heavy metal-resistant bacteria were enumerated from landfill leachate samples (AB-1, MT-1) using the spread-plate method on nutrient agar supplemented with 100 μg/mL individual metals (Pb, Cr, Cd, Cu). Total viable bacterial counts were simultaneously determined on metal-free nutrient agar from identical serial dilutions, yielding AB-1: 4.5×10⁵ CFU/mL and MT-1: 6.2×10⁵ CFU/mL (within typical landfill leachate ranges) [21]. Pb-resistant bacteria showed highest prevalence: AB-1 (0.71% = 3.2×10³/4.5×10⁵ CFU/mL), MT-1 (1.77% = 1.1×10⁴/6.2×10⁵ CFU/mL), followed by Cr-resistant (AB-1: 0.49%; MT-1: 0.97%), Cu-resistant (AB-1: 0.11%; MT-1: 1.61%), and Cd-resistant (AB-1: 0.11%; MT-1: 0.26%) (**S1 Fig and S2, S4 Tables)**. Pb resistance prevalence was highest among all metals tested, consistent with Pb contamination gradients typical of landfill leachate environments.

Of 81 morphologically distinct isolates selected across all metal plates (Table S3), 31 (38%) exhibited Pb resistance, 21 (26%) Cd resistance, 15 (19%) Cr resistance, and 14 (17%) Cu resistance (**S3 Table)**. These isolates were chosen for subsequent analysis according to their size, colony morphology, and pigmentation (**S2 Fig).** This suggests that higher levels of heavy metal contamination may be associated with increased bacterial resistance.

### Maximum Metal Tolerance Limits (MTLs) of Heavy Metal-Resistant Isolates

The MTLs for the 81 heavy metal-resistant isolates were determined across a concentration gradient of 100-3200 μg/mL for Pb, Cr, Cd, and Cu using Mueller-Hinton agar dilution **(S3 Fig)**. Pb-resistant isolates (n=31) exhibited the highest tolerance, with 19.35% (6/31) growing at 3200 μg/mL Pb, 35.48% (11/31) at 1600 μg/mL, 58.06% (18/31) at 800 μg/mL, 93.55% (29/31) at 400 μg/mL, and 96.77% (30/31) at 200 μg/mL **(S4 Fig)**. Cr-resistant isolates (n=15) showed moderate tolerance, with 26.67% (4/15) growing at 800 μg/mL, 80% (12/15) at 400 μg/mL, and 93.33% (14/15) at 200 μg/mL, but no growth at ≥1600 μg/mL Cr **(S5 Fig)**. Cd-resistant isolates (n=21) tolerated up to 14.81% (3/21) at 1600 μg/mL, 33.33% (7/21) at 800 μg/mL, 70.37% (15/21) at 400 μg/mL, and 77.78% (16/21) at 200 μg/mL, with complete inhibition at 3200 μg/mL **(S6 Fig)**. Cu-resistant isolates (n=14) demonstrated 100% growth (14/14) at 200 μg/mL, 64.29% (9/14) at 400 μg/mL, and 21.43% (3/14) at 800 μg/mL, without growth at concentrations ≥1600 μg/mL Cu **(S7 Fig)**. Pb-tolerant isolates consistently showed the highest MTLs across all concentration levels, followed by Cd > Cr > Cu, reflecting differential selective pressures from landfill leachate contamination profiles.

### Determination of Multiple Heavy Metal Resistance in Bacterial Isolates

This research identified multi-heavy metal resistance (MHMR) profiles across 81 bacterial isolates derived from landfill leachate. Using nutrient agar enriched with lead (Pb), cadmium (Cd), chromium (Cr), and copper (Cu) (**S8 Fig)**, 41.98% isolates (34/81) showed resistance to all four metals (**S9 Fig) (S5 Table)**.

### Screening for Plastic-Degradation Potential in Heavy Metal-Resistant Isolates

Additionally, 44.44% bacterial isolates (36/81) exhibited resistance (**S11 Fig)** to multiple heavy metals and plastic degrading capacity as well, including polyethylene (PE), indicating their significant potential for plastic degradation due to their ability to tolerate both environmental pollutants and plastic substrates **(S10 Fig)**.

### Correlation Analysis of Plastic Degradation, Multi-metal Resistance, and Maximum Heavy Metal Tolerance Limits in Bacterial Isolates

Pearson’s correlation analysis was conducted among plastic-degrading isolates, multiple metal-resistant bacteria (bacteria resistant to 4 heavy metals), and maximum metal tolerance limit (MTL) across all 81 isolates **(S6 Table)**. Significant positive correlations were observed between (i) plastic-degrading isolates and multiple metal-resistant bacteria (r = 0.246, p = 0.027) and (ii) maximum metal tolerance limits (MTLs) and multiple metal-resistant bacteria (r = 0.223, p = 0.045), indicating moderate associations between these traits. The correlation between plastic degradation and MTL was weaker and non-significant (r = 0.200, p = 0.074).

### Enzymatic Tests of the Bacterial Isolates

Of the 81 isolates, 25.9% (21) (**S12 Fig)** were urease-positive. The oxidase test showed that 26% (21) (**S13 Fig)** of the isolates demonstrated cytochrome c, as indicated by a purple color change.

All isolates (100%) showed catalase activity (**S14 Fig),** confirmed by rapid bubble formation. The citrate test revealed 18% isolates (11) as citrate-positive (**S15 Fig)**, indicated by a blue color. Esterase activity was assessed in 5 isolates, and 40% (2) exhibited positive activity (**S16 Fig)**, indicated by a halo zone around the colony (**S7 Table).**

### Molecular Characterization of Heavy-metal Resistance and Plastic Degrading and Class 1 Integron Genes

Molecular detection of genetic determinants for lead resistance, plastic degradation, and class 1 integron was conducted on selected isolates. Thirty-one lead-resistant isolates were screened using PCR with the *pbrA*-specific primer (281 bp), and 5 isolates (16%) exhibited a prominent 281 bp amplicon **(Fig 3(a))**. Of the 36 plastic-degrading isolates, 12 were analyzed for the *alkB* gene using a 330 bp primer, with 2 out of 12 isolates (16.7%) showing a 330 bp amplicon **(Fig 3(b))**, indicating their plastic-degrading potential. Additionally, 5 isolates were tested for the class 1 integron gene using the 146 bp primer (intI1), with 4 isolates (80%) exhibiting a prominent 146 bp amplicon **(Fig 3(c))**, confirming the presence of the integron gene.

**Fig 3:**
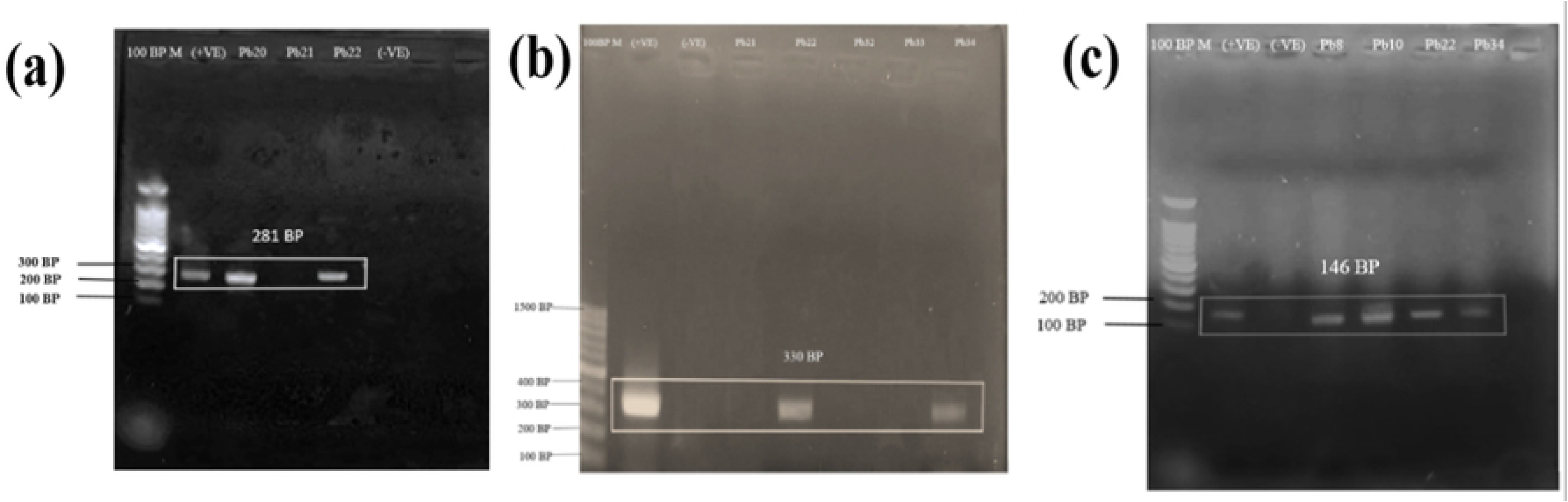
PCR amplicon analysis via 1% agarose gel electrophoresis of **(a)** *pbrA* gene; Lane 1: 100 bp marker (Promega, USA), Lane 2: Positive control, Lane 3: MT-Pb20, Lane 4: MT-Pb21, Lane 5: MT-Pb22 and Lane 6: Negative Control; contain lead resistant isolates containing *pbrA* gene, **(b)** *alkB* gene; Lane 1: 100 bp marker (Promega, USA), Lane 2: Positive control, Lane 3: Negative Control, Lane 4: MT-Pb21, Lane 5: MT-Pb22, Lane 6: MT-Pb32, Lane 7: MT-Pb33 and Lane 8: AB-Pb34; contain lead resistant isolates containing pbrA gene and **(c)** *intI1* Gene. Lane 1: 100 bp marker (Promega, USA), Lane 2: Positive control, Lane 3: MT-Pb20, Lane 4: MT-Pb22, Lane 5: AB-Pb34, Lane 6: AB-Pb8, Lane 7: AB-Pb10 and Lane 8: Negative Control; exhibit the presence of intI1 gene.

### MALDI-TOF MS

MALDI-TOF MS of three heavy metal-resistant isolates identified the MT-Pb20 as ***Bacillus subtilis*,** whereas MT Pb 22 (***Bacillus xiamenensis*)** and AB-Pb 34 (***Bacillus altitudinis***) contains both heavy metal-resistant and plastic-degrading gene. The MALDI spectra for these organisms displayed unique protein peaks that corresponded to those found in the MALDI reference database, providing strong evidence for their identification.

### FTIR Analysis of Metal Removal Capacity

FTIR spectroscopy was employed to characterize the functional groups and identify the chemical bonds involved in the biosorption of lead in MT-Pb20 (***Bacillus subtilis*),** MT-Pb22 **(*Bacillus xiamenensis*)** and AB-Pb34 (***Bacillus altitudinis***). The FTIR spectra of both lead-treated and untreated biomass revealed absorption bands at various wavelengths, indicating the presence of functional groups such as amines (–NH₂), hydroxyl groups (both bonded and free O–H), amides (–CONH–), carboxyl (–COOH), aromatic (C=C) rings, alkenes (C=C), and aliphatic (–CH₂) groups **(Fig 4)**. Differences in peak intensities were observed between the treated and untreated biomass, accompanied by minor shifts in the positions of certain peaks.

**Fig 4:**
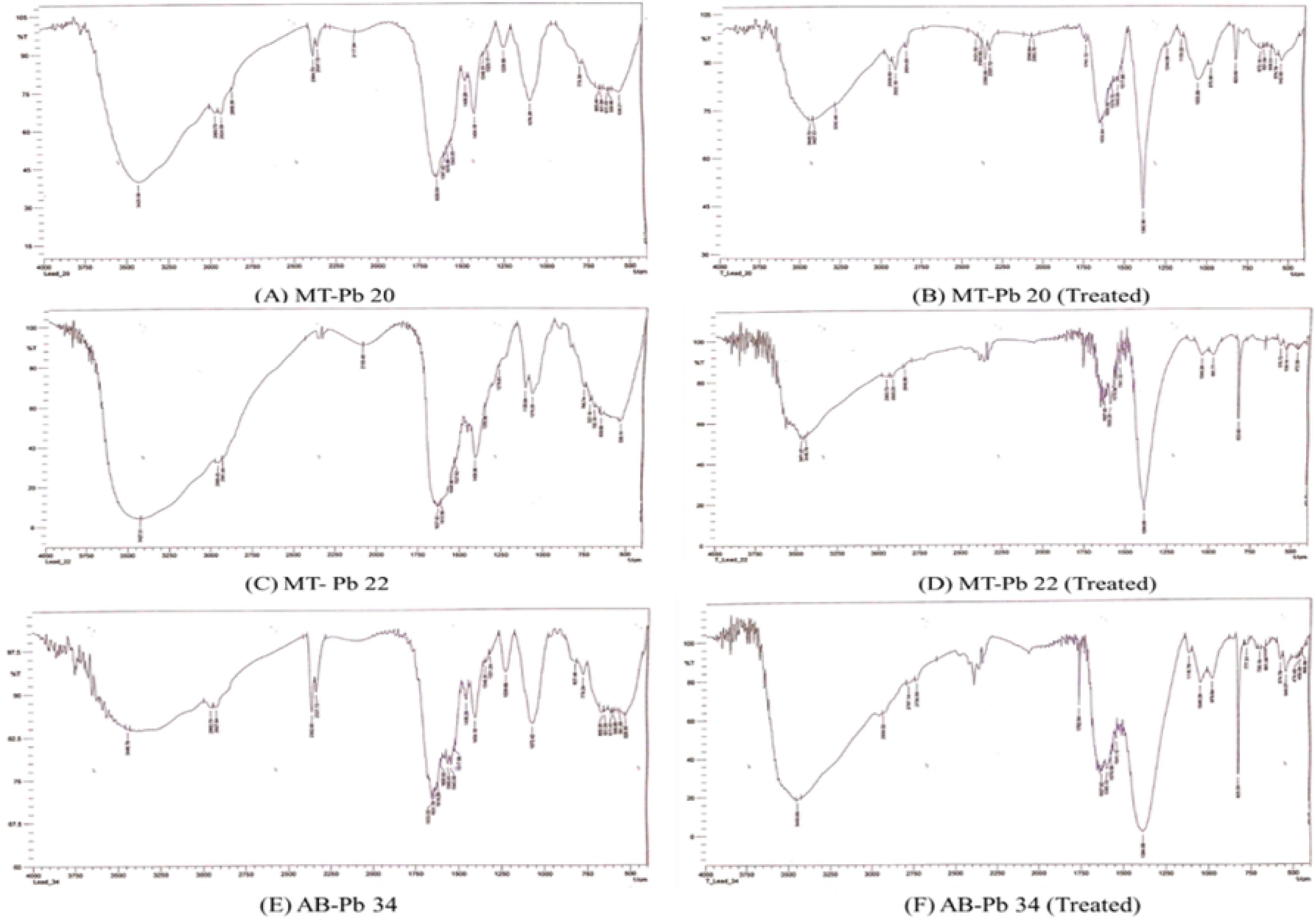
FTIR analysis of untreated biomass (A) MT-Pb20, (C) MT-Pb22 and (E) AB-Pb34 and treated biomass (B) MT-Pb20, (D) MT-Pb22 and (F) AB-Pb34.

## Discussion

Landfill leachate pollution is a significant environmental concern in Bangladesh, particularly at major sites like Aminbazar and Matuail in Dhaka. Leachate from these sites contains high concentrations of heavy metals, organic pollutants, microplastics, and microbial contaminants, which impact microbial ecosystems[26],[27]. Improper leachate treatment poses risks to groundwater and surface water resources [28]. The accumulation of heavy metals and plastics has become a major environmental and public health threat, altering microbial communities [29]. This highlights the urgent need for sustainable leachate management strategies, such as bioremediation. This study aims to characterize heavy metal-resistant and plastic-degrading bacteria from Dhaka’s landfill leachate and evaluate their bioremediation potential.

Leachate samples were collected from various sources, including raw leachate from drainage outlets, untreated, treated, and discharged leachate from a designated pond, and also groundwater. This multi-source sampling approach provides a comprehensive characterization of leachate contamination and its environmental consequences. Data collection was supplemented by a structured questionnaire administered to local community members, ensuring the inclusion of local knowledge and conditions.

Landfills in and around Dhaka support diverse microbial communities that are enriched with metal- and plastic-resistant bacteria, and their persistence appears to be modulated by a suite of physico-chemical parameters typical of highly contaminated landfill leachate environments [30]. Temperature in the contaminated water of Aminbazar and Matuail landfills ranged from 27°C to 32°C and 29.5°C to 37.9°C, respectively, which lies within the mesophilic to thermophilic range favoring the growth and metabolic activity of many bacteria, including those involved in organic and plastic degradation. Consistent with these conditions, thermal profiles in non-sanitary landfills elsewhere in the Indian subcontinent have reached up to about 51°C, generating a distinct microclimate that can select for heat-tolerant and stress-adapted microbial strains equipped to withstand metal and polymer-based pollutants [31].

The pH in this study varied from 6.30 to 9.76, somewhat higher than previously reported values for Aminbazar and Matuail raw leachate (approximately 7.87–8.07) but still within the common range for stabilized landfill leachate (around pH 7–8.5), where alkalinity from bicarbonate and organic acids buffers the system and inhibits the proliferation of acid-sensitive taxa [32]. More basic pH at Matuail is consistent with an advanced methanogenic phase, where acetogenic and methanogenic consortia consume organic acids and elevate pH, thereby reducing metals’ solubility in some cases while favoring alkaliphilic microorganisms adapted to high-pH, low-oxygen conditions [5].

Total dissolved solids (TDS) in raw leachate were elevated, ranging from 120 to 5900 mg/L for Aminbazar and Matuail, closely resembling prior findings for Matuail (662–4650 mg/L) and highlighting the long-standing salinization of leachate from poorly engineered landfills in Dhaka [5]. Corresponding electrical conductivity (EC) values—9061.54 µS/cm in Aminbazar and 4307.69 µS/cm in Matuail—are broadly comparable to a reported 8827 µS/cm for Aminbazar raw leachate, underscoring strong mineralization and ion loading in these waters [4]. The measured salinity of 2.10% in raw leachate from Aminbazar is higher than the previously reported 2.06%, indicating progressive accumulation of dissolved salts in an already highly contaminated landfill environment [4]. Dissolved oxygen (DO) in raw leachate ranged from 0.4 mg/L to 1.07 mg/L, closely matching earlier measurements from Aminbazar and other similar non-sanitary sites where DO typically ranges from 0.87 to 1.33 mg/L and reflects strongly anoxic conditions [4]. Raw leachates exhibited elevated pH, TDS, EC, and salinity alongside lower DO relative to treated or receiving-water samples, consistent with the highly mineralized and low-oxygen leachate characteristics described for poorly engineered landfills, which makes them appropriate matrices for downstream microbial and ecotoxicological analysis [33].

Heavy metal concentrations in raw landfill leachate significantly exceed WHO permissible limits of 0.05 mg/L for Pb, 0.005 mg/L for Cd, 0.05 mg/L for Cr, and 0.05 mg/L for Cu [34]. Pb concentrations ranged from 16.6 to 30.32 mg/L, Cr from 14.06 to 24.58 mg/L, Cu from 5.47 to 21.05 mg/L, and Cd from 0.69 to 0.88 mg/L, all substantially higher than those reported in other studies [35],[36]. These elevated levels indicate severe contamination, posing significant environmental and health risks.

Bacterial isolates from landfill leachate exhibited varying levels of heavy metal tolerance. Lead-resistant isolates ranged from 32×10² to 110×10² cfu/ml, lower than the 54×10⁴ cfu/ml found in tannery effluents [37]. Chromium-resistant isolates ranged from 22×10² to 60×10² cfu/ml, with no similar counts reported in other leachate studies. Cadmium-resistant isolates ranged from 5×10² to 16×10² cfu/ml, lower than the 7×10⁴ cfu/ml reported by Renu et al. (2022) ^34^. A significant Pearson correlation (p < 0.05) between AAS-measured heavy metals and total bacterial count suggests that higher heavy metal contamination may be associated with increased bacterial resistance. A total of 81 isolates, including 31 Pb-resistant, 15 Cr-resistant, 21 Cd-resistant, and 14 Cu-resistant isolates, were selected for further experiments based on their colony morphology, size, and resistance characteristics.

MTL assays revealed varying tolerance levels in metal-tolerant bacterial isolates. Pb-resistant strains tolerated up to 3200 μg/mL, with 96.77% (30/31) of isolates withstanding 200 μg/mL Pb, higher than the 1950 μg/mL tolerance reported for Gemella sp. [37]. Cr, Cd, and Cu resistance varied, with some isolates showing no growth above 800 μg/mL. Four Cd-resistant isolates tolerated up to 1600 μg/mL, and four Cr-resistant and three Cu-resistant isolates tolerated up to 800 μg/mL, consistent with previous studies [37]. These findings suggest the potential use of these isolates in bioremediation of heavily contaminated sites, such as landfills.

The study examined bacterial resistance to multiple heavy metals and plastic degradation. Among 81 isolates, 42% (34) were resistant to Pb, Cr, Cd, and Cu, while 36% (29), 18% (15), and 4% (3) were resistant to three, two, and one metal, respectively, indicating the selective pressure of heavy metal contamination in landfills. Additionally, 44% (36) of the isolates exhibited resistance to plastic materials like polyethylene, suggesting potential for sustainable waste management and bioremediation. A significant positive correlation was observed between plastic degradation capacity and multimetal resistance (Pb, Cr, Cd, Cu) (r = 0.246, p = 0.027), suggesting potential shared adaptive mechanisms such as efflux pumps or oxidative stress responses that confer tolerance to both plastic-derived oligomers and heavy metals [38]. Similarly, maximum metal tolerance limits (MTLs) positively correlated with multimetal resistance (r = 0.223, p = 0.045), consistent with reports of co-occurring resistance traits in polluted environments where bacteria evolve broad stress tolerance. The weaker, non-significant correlation between plastic degradation and MTLs (r = 0.200, p = 0.074) indicates these traits may involve partially distinct pathways, as plastic degradation often relies on specialized hydrolases while metal tolerance depends on sequestration and extrusion systems a pattern seen in *Pseudomonas* and *Bacillus* isolates from contaminated sites. Other previous studies, such as Keramati et al. (2011) [39] and Di Cesare et al. (2021) [40], found similar findings in *Pseudomonas* and *Rhodococcus ruber*, which degrade plastics while exhibiting resistance to heavy metals and antibiotics in some cases, highlighting their potential for bioremediation [18,39].

Enzymatic profiles of bacterial isolates revealed their metabolic capabilities, particularly in nitrogen cycling, oxidative stress response, and environmental adaptability. In the urease test, 60% of isolates were urease-positive, consistent with Margesin et al. (1999)[41], suggesting a role in Microbially Induced Calcite Precipitation (MICP) for pollutant immobilization[25]. However, 40% were urease-negative, contrasting with Karigar and Rao (2011) [42], who reported higher urease activity. In the oxidase test, 26% of isolates were oxidase-positive, indicating adaptation to low-oxygen environments [43]. All isolates were catalase-positive, aligning with, demonstrating their ability to neutralize reactive oxygen species (ROS) in oxidative stress conditions [44]. Only 18% were citrate-positive, consistent with Karigar and Rao (2011) [42], suggesting that citrate is not a preferred carbon source in these bacteria. Additionally, isolates containing the *alkB* gene showed positive esterase activity. These profiles suggest that the isolates are metabolically versatile, with potential for bioremediation in contaminated environments [45].

PCR detection of *pbrA* (lead resistance gene), *alkB* (plastic-degrading gene), and *int1* (class 1 integron genes) confirmed the genetic basis of resistance in selected isolates. The presence of integron genes suggests potential horizontal gene transfer (HGT) of resistance traits within microbial communities [46]. This supports the hypothesis that landfill leachates may serve as reservoirs of resistant bacteria, influencing microbial diversity in surrounding ecosystems. Exposure to metals can increase the prevalence of heavy metal resistance genes (HMRGs) in bacteria [47]. In this study, 16% (5/31) of Pb-resistant isolates showed the presence of the *pbrA* gene. Among these 5 HMRG-containing isolates, 40% (2/5) also contained the *alkB* gene. While this study prioritizes the *alkB* gene, the landfill microbiome harbors diverse plastic-degrading genes, including laccase and alkane monooxygenases (*alkM*, *alkB1_2*, *ladA*) [2]. Class 1 integrons, which capture and express gene cassettes linked to resistance, were found in 80% (4/5) of Pb-resistant isolates. That means these four isolates also exhibited HMRGs, and two (MT-Pb22 and AB-Pb34) also contained the *alkB* gene, indicating potential for both metal tolerance and plastic degradation. These findings align with the previous studies in Bangladesh and elsewhere[48,49] highlighting the bioremediation potential of these isolates in polluted environments.

Exposure to heavy metals has led to the development of resistance mechanisms in microbial habitats, which can be used for the detoxification and degradation of these pollutants. In this study, three isolates - MT-Pb20 (*Bacillus subtilis)*, MT-Pb22 (*Bacillus xiamenensis*), and AB-Pb34 (*Bacillus altitudinis*) identified by MALDI-TOF MS demonstrated significant removal of Pb, confirmed by FTIR analysis, highlighting their potential for bioremediation of heavy metal contamination. Previous studies have reported *Pseudomonas sp.*, *Klebsiella pneumonia*, and *Stenotrophomonas maltophila* as effective bioremediators [49]. Several bioremediation processes, including biosorption, bioleaching, biomineralization, biotransformation, and intracellular accumulation, are employed for the effective removal of heavy metals **(Fig 5)** [54]. FTIR spectroscopy revealed functional group interactions (hydroxyl -OH, carboxyl -COOH, and carbonyl C=O) in untreated biomass, with shifts in peak intensities in treated biomass after metal biosorption **(Table 2)**, indicating successful metal binding [55].

**Fig 5:**
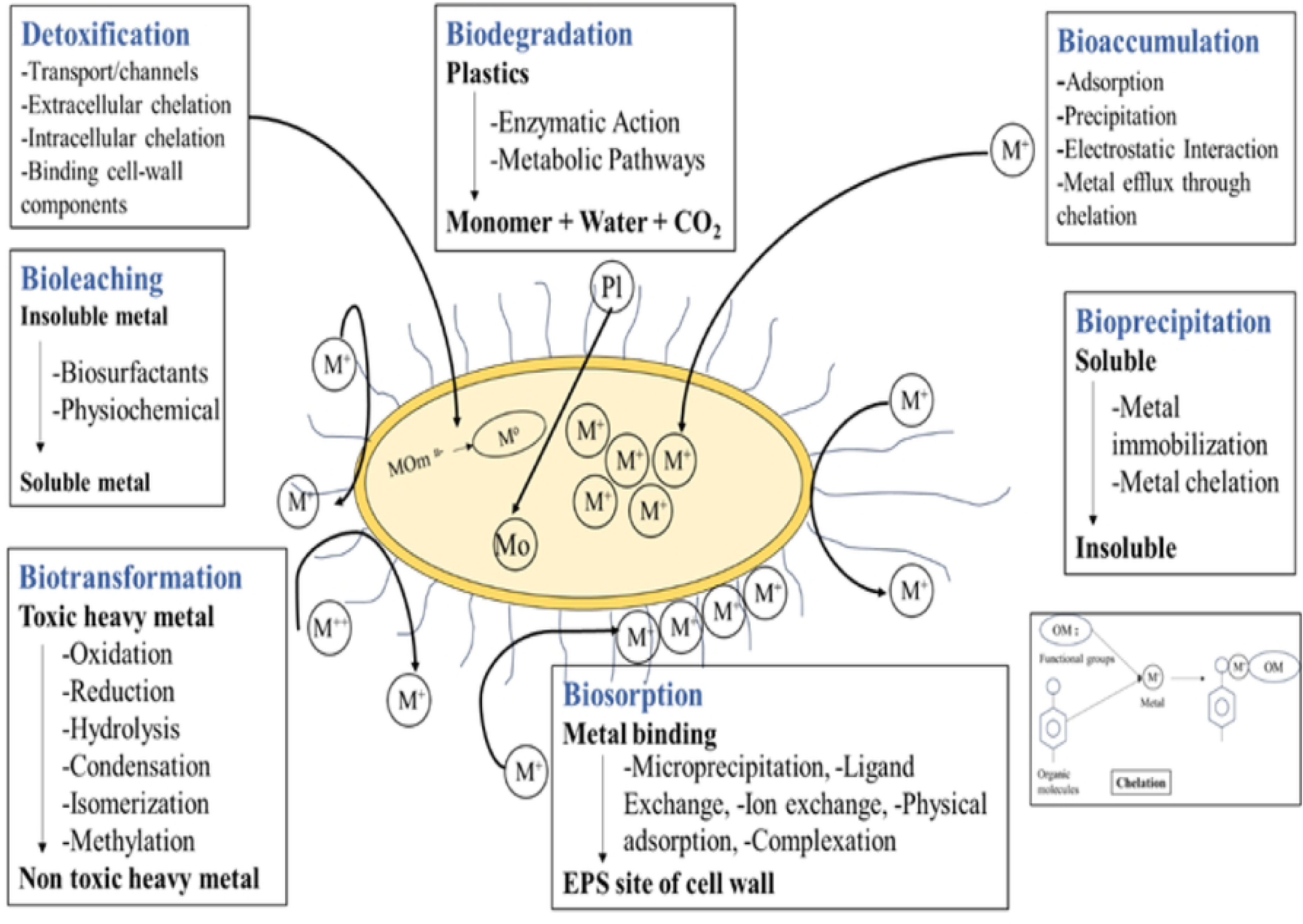
Mechanism of Heavy-metal and Plastic Bioremediation.

**Table 2.**
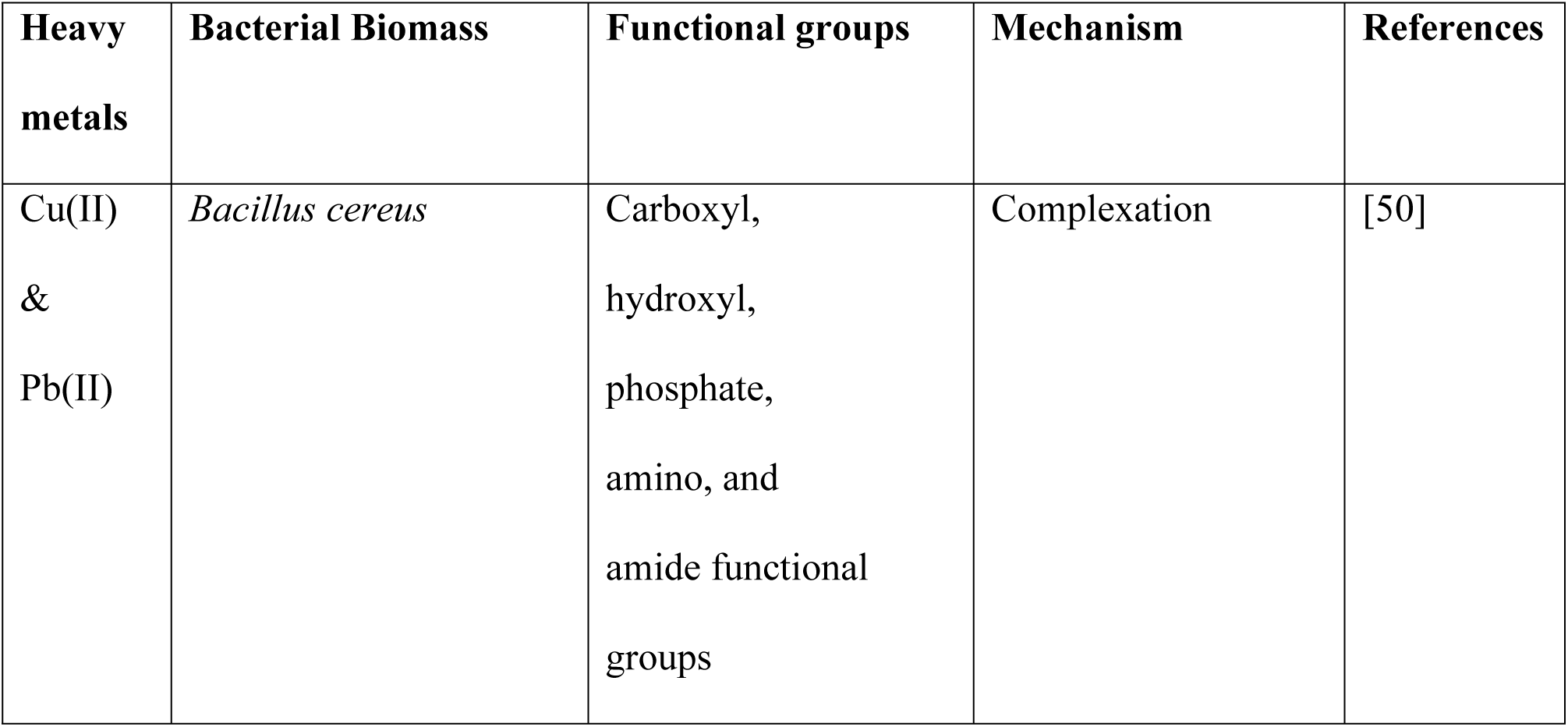
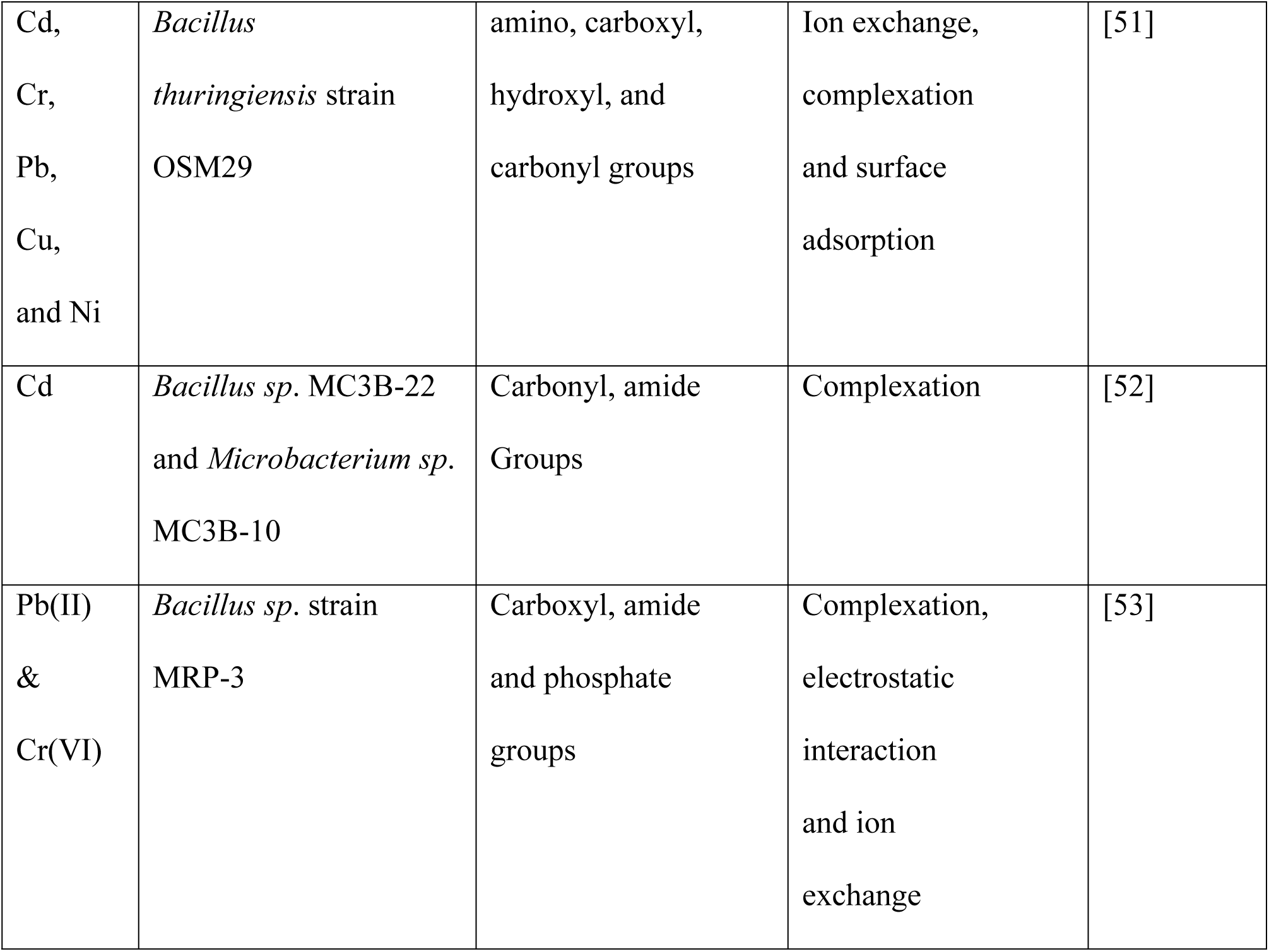
Different functional groups of bacteria that interact with heavy metals.

| Heavy metals | Bacterial Biomass | Functional groups | Mechanism | References |
| --- | --- | --- | --- | --- |
| Cu(II)<br>&<br>Pb(II) | *Bacillus cereus* | Carboxyl,<br>hydroxyl,<br>phosphate,<br>amino, and<br>amide functional<br>groups | Complexation | [50] |
| Cd,<br>Cr,<br>Pb,<br>Cu,<br>and Ni | *Bacillus thuringiensis* strain OSM29 | amino, carboxyl, hydroxyl, and carbonyl groups | Ion exchange, complexation and surface adsorption | [51] |
| Cd | *Bacillus sp.* MC3B-22 and *Microbacterium sp.* MC3B-10 | Carbonyl, amide Groups | Complexation | [52] |
| Pb(II) & Cr(VI) | *Bacillus sp.* strain MRP-3 | Carboxyl, amide and phosphate groups | Complexation, electrostatic interaction and ion exchange | [53] |

A strong absorption band at 3400 cm⁻¹ in the untreated biomass corresponded to N–H stretching of amines or O–H stretching of hydroxyl groups, either free or hydrogen-bonded in untreated ***Bacillus subtilis*, *Bacillus xiamenensis*** and ***Bacillus altitudinis***. In contrast, the lead-exposed biomass displayed a broad vibrational band ranging from 3200 to 3600 cm⁻¹ without a sharp peak, likely due to overlapping of amine and hydroxyl group stretching or interactions between lead ions and these functional groups. In both lead-exposed and unexposed biomass in ***Bacillus subtilis*** [56], the absorption peak at 2925 cm⁻¹ corresponding to aliphatic (–CH₂) groups remained unchanged. This suggests the presence of asymmetric stretching vibrations associated with proteins, lipids, nucleic acids, and polysaccharides within the biomass cell wall [57]. It displayed minor shifts in FTIR absorption peaks from 2364.73 to 2366.66, 2117.84 to 2065.76, 1587.42 to 1600.92, 1228.66 to 1244.09, 1076.28 to 1055.06 and a major shift from 1404.18 to 1382.96 in Pb treated and untreated biomass suggesting interaction of lead with functional groups on the biomass surface [58].

In ***Bacillus xiamenensis***, the absorption peak at 1627.92 cm⁻¹ corresponding to the amide I (C=O) group of proteins remained unchanged. The FTIR spectrum of Pb-exposed biomass revealed notable shifts in absorption peaks, particularly from 1409.96 to 1384.89 cm⁻¹ and 1076.28 to 1049.28 cm⁻¹, indicating the involvement of carboxyl and C–O/P–O functional groups in Pb binding [59]. Additional minor shifts from 3427.51 to 3446.79 cm⁻¹ in the O–H/N–H stretching region and from 2968.45 to 2960.73 cm⁻¹ in the C–H stretching region further support alterations in the biomass surface chemistry due to Pb interaction.

In ***Bacillus altitudinis*** [60], the FTIR spectrum of Pb-exposed biomass reevealed notable shifts in absorption peaks, particularly from 1409.96 to 1384.89 cm⁻¹ and 1076.28 to 1049.28 cm⁻¹, indicating the involvement of carboxyl and C–O/P–O functional groups in Pb binding. Minor shifts from 3446.79 to 3450.65 cm⁻¹ (O–H/N–H), 2960.73 to 2939.52 cm⁻¹ (aliphatic C–H), and 1653.00 to 1627.92 cm⁻¹ (amide I) further support changes in biomass surface chemistry. The shift from 2362.80 to 2340.99 cm⁻¹ may reflect weakly bound groups or atmospheric interference [59].

Therefore, the FTIR analysis of Pb-exposed bacterial biomass indicated that the observed changes were likely caused by interactions between lead and organic functional groups such as amino, hydroxyl, and carboxyl groups through chelation on the biomass cell surface.

### Conclusions

This study examined bacterial communities from landfill leachate in Dhaka, Bangladesh, focusing on heavy metal resistance and plastic degradation. Out of 81 bacterial isolates, 42% exhibited resistance to Pb, Cr, Cd, and Cu, with Pb showing the highest resistance (up to 3200 μg/mL). Notably, 44% of the isolates demonstrated plastic degradation abilities. The study found that 25.9% of isolates were urease-positive which has potential role in bioremediation. Genetic analysis revealed the presence of metal resistance genes *pbrA* for Pb, and some isolates also contained the *alkB* gene for plastic degradation. MALDI-TOF MS identified ***Bacillus subtilis*, *Bacillus xiamenensis*** and ***Bacillus altitudinis*** showing significant bioremediation capability also, marking it as the first of its kind reported in Bangladesh.

## Methods and Materials

### Sampling, Transportation, and Physicochemical Analysis

The study, conducted between June 2023 to June 2024, covered the two primary landfills serving Dhaka city are the Aminbazar landfill, located in the northern region near the Dhaka-Aricha Highway (23°48’03’N, 90°18’30’E), and the Matuail landfill, located in the southern region approximately 8 kilometers south of Gulistan (23°42’23’N, 90°25’10’E), both of which manage large share of the municipal solid waste produced by Dhaka North City Corporation (DNCC) and Dhaka South City Corporation (DSCC) respectively **(Fig 6)** [4].

**Fig 6:**
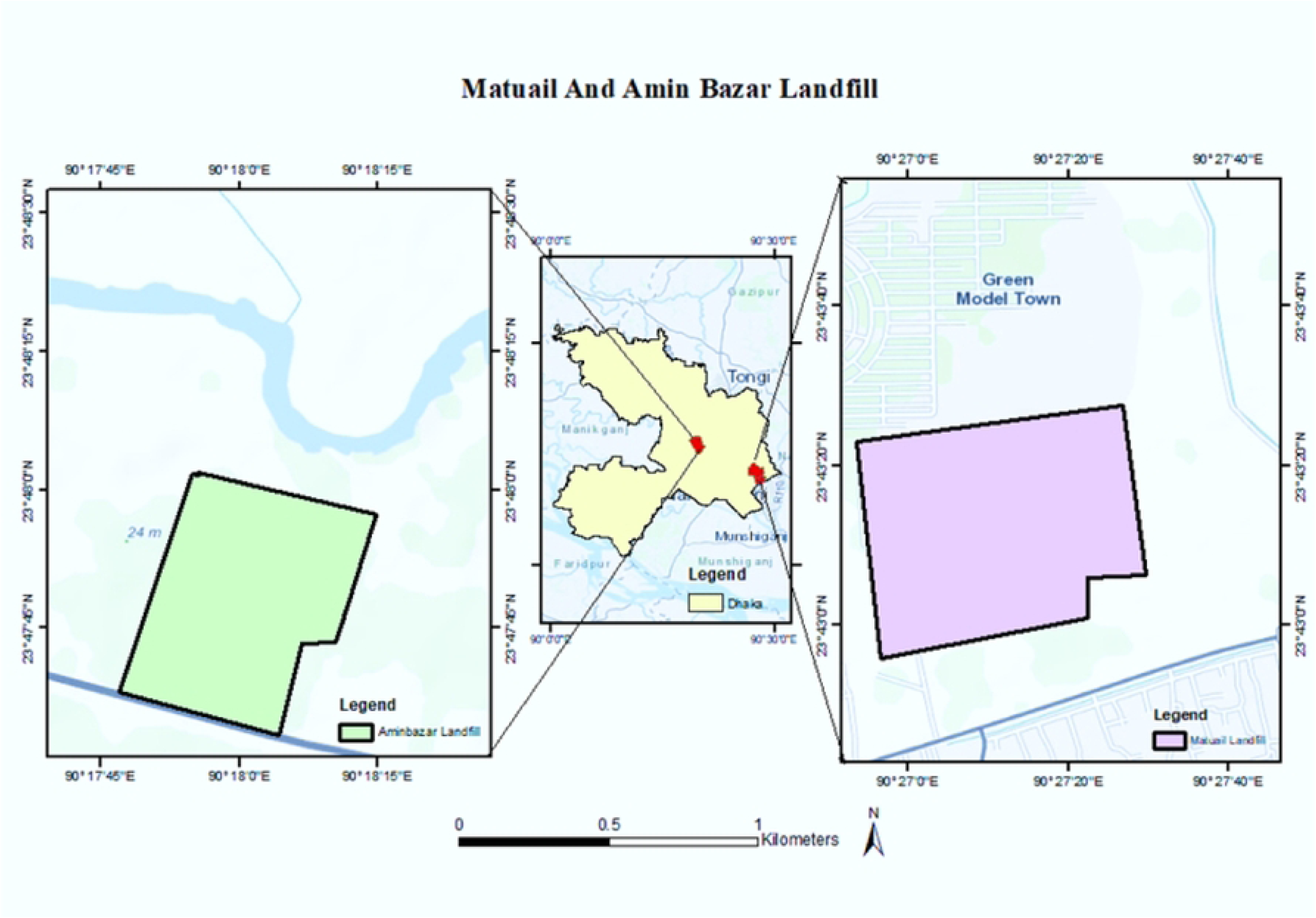
Geographical Representation of the Sampling Site.

A total of 10 samples, comprising 8 leachate and 2 groundwater samples, were systematically collected. Meanwhile, water samples were placed in 500 ml sterile Duran bottles (Schott, Germany), sealed tightly, and transported to the lab in insulated ice boxes for quick bacteriological analysis. Physicochemical parameters, including pH, temperature, total dissolved solids (TDS), electrical conductivity (EC), salinity, and dissolved oxygen (DO), were quantified using a thermometer, pH meter electrode (Orion-2 STAR, Thermoscientific), and a DO meter (970 DO2 meter, Jenway UK).

### Analysis of Metal Concentration in the Leachate Samples

Heavy metal concentrations (Pb, Cr, Cd, and Cu) were determined using the acid digestion method. Leachate samples (300 mL) were dried at 105°C, pulverized, and subsequently digested with a 4:1 mixture of HNO₃ and HClO₄ at 450°C for a duration of 1 hour. The resulting digests were filtered through Whatman 541 paper, diluted to 25 mL with distilled water, and analyzed using an Atomic Absorption Spectrophotometer (Shimadzu, AA-6701F) [61]. Calibration curves were generated using standard solutions were prepared by diluting 1000 mg/L stock solutions, and concentrations were determined using the air-acetylene flame technique (APHA 3111-B) [62].

### Primary Screening for Heavy Metal-resistant Bacteria

Heavy metal-resistant bacteria were screened using nutrient agar (Scharlau, Spain) as the base medium, which was supplemented with 100 μg/ml metal solutions prepared by diluting 1000 mg/L stock solutions. Stock solutions of lead (Pb), chromium (Cr), cadmium (Cd), and copper (Cu) were prepared in Milli-Q water from analytical-grade Pb(NO₃)₂, K₂Cr₂O₇, 3CdSO₄·8H₂O, and CuSO₄·5H₂O, and subsequently sterilized by filtration through 0.22 μm membrane filters to obtain a final concentration of 1000 mg/L [63]. A 1 ml sample of collected landfill leachate was mixed with 9 ml of normal saline (0.85%) and serially diluted from 10⁻¹ to 10⁻⁴. From the 10⁻², 10⁻³, and 10⁻⁴ dilutions, 100 μl samples were spread onto metal-supplemented nutrient agar plates, which were then sealed. The plates were incubated at 37± 1°C for a duration of 24 to 72 hours [64].

Following incubation, the plates were examined for distinct bacterial colonies exhibiting variations in morphology, including shape, elevation, margin, surface texture, and size. Bacterial growth was quantified as colony-forming units (cfu) per milliliter, and individual representative colonies were selected and isolated through streak plate cultivation to yield pure cultures. The purified and well-grown colonies were preserved at −20°C.

### Assessment of the Maximum Metal Tolerance Limits (MTLs) of Heavy Metals

The maximum tolerance limit (MTL) for each heavy metal was determined following the approach described by Abbas et al. (2025) in *Frontiers in Microbiology*. In this method, the MTL is defined as the highest metal concentration at which visible bacterial growth is still observed, whereas complete inhibition of growth at the next higher concentration is considered the threshold [65]. The MLTs for metal-tolerant strains was determined using Mueller–Hinton agar (HIMEDIA, India), supplemented with lead, cadmium, chromium, and copper at final concentrations ranging from 100 to 3200 μg/mL [66,67]. Metal salt stock solutions were prepared in Milli-Q water, sterilized by 0.22 μm filtration, and incorporated into MH agar at the desired final concentrations. Plates included growth controls (MH agar without metal) and sterility controls (metal-supplemented MH agar without inoculum) also. The heavy metal-supplemented plates were divided into four equal sectors, and each isolate was streaked in a separate quadrant. All assays were conducted in triplicate. Plates were incubated at 37± 1°C for 4–6 days, after which bacterial growth was assessed and the MTL was recorded as the highest metal concentration at which the tested isolate no longer exhibited growth [65].

### Multi-metal Resistance Capacity

A multi-metal resistance test was conducted on each of the individual heavy metal-resistant isolates to assess their capacity to tolerate additional heavy metals, such as lead (Pb), chromium (Cr), cadmium (Cd), and copper (Cu). The isolates were streaked onto nutrient agar (NA) plates supplemented with 100 μg/mL of lead nitrate (Pb(NO₃)₂), cadmium sulfate (CdSO₄·8H₂O), potassium dichromate (K₂Cr₂O₇), and copper sulfate (CuSO₄). The plates were incubated at 37± 1°C for 24, 48, and 72 hours, following which the isolates were assessed for their resistance to these metals [68]. The growth patterns observed provided valuable data on the multi-metal resistance characteristics of the isolates.

### Detection of Plastic Degrading Isolates

To ensure the isolated microorganisms were capable of degrading plastic, a minimal salt medium (MSM) was prepared containing Na₂HPO₄ (6.0 g/L), KH₂PO₄ (3.0 g/L), NH₄Cl (1.0 g/L), NaCl (0.5 g/L), MgSO₄·7H₂O (0.1 g/L), FeSO₄·7H₂O (0.01 g/L), and trace elements such as ZnSO₄ and MnCl₂. Agar (1.5% w/v) was incorporated into the medium to solidify it for petri plate preparation. 1 g of polyethylene powder (low density, 500 microns) (Cat: A10239.22, United Kingdom), used as the sole carbon source, it was incorporated into the medium. The isolated microorganisms were then inoculated into MSM plates containing the polyethylene, the plates were sealed (parafilm) to prevent desiccation and contamination, and incubated at 30°C for 30 days. A control experiment, without bacterial inoculation, was conducted under the same conditions to differentiate between biotic and abiotic degradation. This method effectively identified able to use polyethylene as a carbon source, demonstrating their plastic-degrading potential [26].

### Enzymatic Assay Methods

The urease test detects urease activity, which catalyzes the hydrolysis of urea into ammonia (NH₃) and carbon dioxide (CO₂), leading to an increase in pH. A positive result is characterized by a color change to pink (pH 8.1), indicating an increase in pH, as observed after 24 hours [69]. The oxidase test identifies the presence of cytochrome c oxidase, an enzyme involved in the electron transport chain. A positive result is marked by the appearance of a dark purple color on a filter paper soaked with tetramethyl-p-phenylenediamine dihydrochloride. The catalase test detects catalase activity, where hydrogen peroxide (H₂O₂) is broken down into water and oxygen, forming visible bubbles. The immediate appearance of bubbles after adding hydrogen peroxide indicates a positive result [70]. The esterase test identifies esterase activity, which catalyzes the hydrolysis of ester bonds in organic compounds, including pesticides. Positive activity is indicated by the formation of a clear halo around colonies on Tween 80 agar, incubated at 28°C for 5 days [24].

### Molecular Characterization of Heavy Metal (Lead) Resistance and Plastic Degrading Genes

Bacterial genomic DNA was extracted using a boiling method. PCR was conducted to identify resistance gene patterns. The reaction mixture consisted of a total volume of 25 μl, including 12.5 μl of master mix, 1 μl each of forward (10 pmol) and reverse primers (10 pmol), 6.5 μl of nuclease-free water, and 4 μl of DNA template (**S6 Table).**

PCR amplification of lead resistance gene, *pbrA* (primers: F-CCTCGCCATCGATCACTACC, R-GCACCAGTGCATCACGAATC) was performed at 94°C for 2 min, followed by 40 cycles consisting of denaturation at 94°C for 1 min, annealing at 60°C for 1 min, extension at 72 °C for 3 min and a final extension of 72 °C for 7 min, resulting in an amplicon of 281 base pairs (bp) [71]. PCR amplification of plastic degrading gene, *alkB* (primers: F-

TCGAGCACAACCGCGGCCACCA, R-CCGTAGTGCTCGACGTAGTT) was conducted at 95°C for 15 min, followed by 39 cycles consisting of denaturation at 94°C for 1 min, annealing at 55°C for 1 min, extension at 72 °C for 1 min and a final extension of 72 °C for 10 min, yielding an amplicon of 330 bp [26]. For detection of Class 1 Integron *intI1* genes (primers: F-GGCTTCGTGATGCCTGCTT, R-CATTCCTGGCCGTGGTTCT), PCR amplification was performed in a thermocycler based on the following programme: initial denaturation at 94°C for 1 minutes, 35 cycles of denaturation at 98°C for 30 seconds, annealing at 58°C for 30 seconds and a final extension at 72°C for 10 min, producing an amplicon of 146 bp [72]. For each primer set, reactions included a positive control (genomic DNA from a reference strain previously confirmed to carry the respective target gene) and a negative control (no-template control) to monitor for non-specific amplification and contamination. The resulting amplified products were then separated by agarose gel electrophoresis on a 1% gel stained with ethidium bromide and examined under a UV transilluminator.

### Identification of Isolates Using MALDI-ToF MS

Heavy metal- and plastic-degrading pure isolates were identified by MALDI-TOF MS at NAHDIC following Toubal et al. (2018) and the validated Bacillus protocol of Starostin et al. (2015) [73] [74]. In this method, twelve colonies per isolate were extracted (ethanol inactivation, 70% formic acid cell wall disruption, acetonitrile protein extraction), spotted in triplicate onto stainless steel plates, overlaid with α-cyano-4-hydroxycinnamic acid matrix, and analyzed on a Bruker Ultraflex III MALDI-TOF/TOF (linear positive mode, 25 kV, 2000–20,000 Da, 500 laser shots/spectrum) calibrated with E. coli ribosomal proteins [75]. Spectra were identified using Biotyper 3.1 with manufacturer thresholds (≥2.0 species-level, 1.7–1.99 genus-level); only reproducible high-confidence identifications across replicates were accepted, achieving 100% accuracy for Bacillus pumilus group strains [74].

### Determination of Metal Removal Capacity Using Fourier Transform Infrared Spectroscopy (FTIR)

FTIR spectroscopy was used to analyze bacterial biomass treated with lead (Pb) and chromium (Cr) to examine structural and functional changes induced by heavy metal stress [76]. The biomass, prepared as diamond disks, was analyzed in the 4000–400 cm⁻¹ range using a UATR Two FTIR spectrometer (Perkin Elmer, USA) at 4 cm⁻¹ resolution, with KBr pellets for background correction. Untreated biomass was also analyzed using the IRPrestige-21 Shimadzu spectrometer (Figure 2.8). FTIR detected changes in functional groups like hydroxyl (-OH), carboxyl (-COOH), and phosphate (-PO₄), which are crucial for metal binding [77].

### Statistical Analysis

Statistical analysis was conducted using Microsoft Excel 2016 (Microsoft Corporation, Redmond, WA, USA) for descriptive statistics and IBM SPSS Statistics Version 23 for inferential analysis. Pearson’s correlation coefficient (r) was computed to evaluate relationships between variables (heavy metal concentrations, minimum tolerated levels, plastic degradation rates), with two-tailed tests at α=0.05 significance. Correlations achieving p<0.05 were considered statistically significant.

### Limitations

However, further studies with larger sample sizes and long-term data are needed to assess contamination levels, microbial responses, and seasonal variations. Additionally, MALDI-ToF MS identifications were not systematically confirmed by 16S rRNA gene sequencing, more comprehensive gene identification is required to understand the genetic mechanisms of metal resistance and plastic degradation, as well as to optimize plastic degradation for identifying bioremediation potential.

## Author’s contribution

Umme Samia Antu: Planned the projects, conducted the experiments, analyzed the data, and wrote the original manuscript; Amily Sarker, Nabila Haque and Joyoshrie Karmakar: Helped during the experiments; Abdul Khaleque: Helped in writing-review and editing; Md. Sabir Hossain and Md. Anowar Khasru Parvez: Conceptualization, Designed and supervised the project, Writing-Review and Editing. All the authors approved the manuscript’s final version for publication.

## Ethics approval

No need to ethical permission

## Data availability

All data generated or analyzed during this study are included in this published article, along with its supplementary information files.

## Abbreviation

AB: Aminbazar
MT: Matuail
AB-1: Aminbazar Raw Leachate
MT-1: Matuail Raw Leachate
DO: Dissolved Oxygen
BOD: Biochemical Oxygen Demand
EC: Electrical Conductivity
MSW: Municipal Solid Waste
AAS: Atomic Absorption Spectroscopy
CFU: Colony Forming Unit
TBC: Total Bacterial Count
PET: Polyethylene Terephthalate
PE: Polyethylene
F: Forward
AMR: Antimicrobial Resistance
R: Reverse
ATCC: American Type Culture Collection
BTS: Bacterial Test Standard
DNA: Deoxyribo Nucleic Acid
E. coli: Escherichia coli
EDTA: Ethilenediamine Tetraacetic Acid
et al: And others
EtBr: Ethidium Bromide
H_2_O_2_: Hydrogen PER Oxide
HGT: Horizontal Gene Transfer
i.e.: That is
ICU: Intensive Care Unit
MDR: Multidrug Resistant
MGE: Mobile Genetic Element
MEGA: Molecular evolutionary genetic analysis
MHA: Muller –Hinton Agar
Mac: MacConkey Agar
MIC: Minimum Inhibitory Concentration
Min: Minute
MR: Methyl Red
NA: Nalidixic Acid
NA: Nutrient Agar
NCBI: National Center for Biotechnology Information
O_2_: Oxygen
PBP: Penicillin Binding Protein
PCR: Polymerase Chain Reaction
pH: Negative Logarithm of Hydrogen Ion
PMQR: Plasmid Mediated Quinolone Resistance
rRNA: Ribosomal RNA
RNA: Ribonucleic Acid
Rmp: Rotation per minute
Spp: Species
Sec: Second
TSA: Trypticase Soy Agar
UK: United Kingdom
USA: United States OF America
UTI: Urinary Tract Infection
VAP: Ventilator – Associated Pneumonia
MICP: Microbially Induced Calcium Carbonate Precipitation
WHO: World Health Organization
XDR: Extensively Drug Resistance
HMRG: Heavy Metal Resistant Gene
PDG: Plastic Degrading Gene
Cu: Copper
CO: Cobalt
Pb: Lead
Cd: Cadmium
FTIR: Fourier-transform infrared spectroscopy
TDS: Total Dissolved Solids
MTL: Maximum Metal Tolerance Limits

## Symbols

| Symbol | Meaning |
| --- | --- |
| % | Percentage |
| < | Less than |
| > | Greater than |
| ≥ | Less than or equal |
| ≤ | Greater than or equal |
| Mg | Microgram |
| μl | Microliter |
| °C | Degree Celsius |
| Bp | Base pair |
| Kb | Kilobase |
| Kbp | Kilobase pair |
| MDa | Megadalton |
| L | Liter |
| mL | Milliliter |

## Acknowledgements

We sincerely thank the Department of Biochemistry and Molecular Biology, Department of Microbiology, Environmental Health and Synthetic Biology Laboratory, and Wazed Miah Science

Research Center at Jahangirnagar University for providing essential facilities, resources, and collaborative support throughout this research.

